# The META tool optimizes metagenomic analyses across sequencing platforms and classifiers

**DOI:** 10.1101/2021.07.29.454031

**Authors:** Robert A. Player, Angeline M. Aguinaldo, Brian B. Merritt, Lisa N. Maszkiewicz, Oluwaferanmi E. Adeyemo, Ellen R. Forsyth, Kathleen J. Verratti, Brant W. Chee, Sarah L. Grady, Christopher E. Bradburne

## Abstract

A major challenge in the field of metagenomics is the selection of the correct combination of sequencing platform and downstream metagenomic analysis algorithm, or ‘classifier’. Here, we present the Metagenomic Evaluation Tool Analyzer (META), which produces simulated data and facilitates platform and algorithm selection for any given metagenomic use case. META-generated *in silico* read data are modular, scalable, and reflect user-defined community profiles, while the downstream analysis is done using a variety of metagenomic classifiers. Reported results include information on resource utilization, time-to-answer, and performance. Real-world data can also be analyzed using selected classifiers and results benchmarked against simulations. To test the utility of the META software, simulated data was compared to real-world viral and bacterial metagenomic samples run on four different sequencers and analyzed using 12 metagenomic classifiers. Lastly, we introduce ‘META Score’: a unified, quantitative value which rates an analytic classifier’s ability to both identify and count taxa in a representative sample.

## INTRODUCTION

Since its inception, the field of metagenomics has proven to be one of the most challenging arenas of genomics research. Microbial communities are often dynamic, and the countless tools available for characterization all present their own strengths and weaknesses. Experimental design choices are frequently made with reagent cost, availability, and protocol ease in mind, with less emphasis placed on a thorough understanding of the limitations of a particular sequencer, metagenomic classification algorithm, reference database, or the complexity of the sample itself. Here, we introduce the Metagenomic Evaluation Tool Analyzer, or ‘META’, as a solution for predicting and testing the best approach for a given metagenomic experiment (Table 1). Users are able to use this modular and easily updateable software to select, simulate, and compare the performance of different sequencer/classifier combinations towards more efficient use of available wetlab resources.

**Table 1.** META features and rationale.

| META feature | Implemented by | Enables | Achieves |
| --- | --- | --- | --- |
| Modular | Dockers | Futureproofing via ease of updating | The latest, best analysis algorithms |
|  |  | Ease of expanding as new classifiers are published |  |
|  |  | Classification tools with diverse dependencies |  |
| Scalable | Choice of tools and database size | Broader range of deployment options based on available hardware | Validated, deployable bioinformatics |
|  |  | Selection of optimal pipeline at large scale, and deployment on laptop |  |
| Simulation | InSilicoSeq and DeepSimulator | Read generation from a single source across sequencers | Downselection of sequencer and analysis |
|  |  | Evaluation and selection of analysis algorithms without costly sequencing |  |
| Visualization | D3.js | Interactive visualizations | Ease of use and interpretation |
|  |  | Comparison of classifiers based on simulated or real reads |  |
| Resource Estimation | Tracking of compute time, RAM usage | Data driven hardware assessment<br>Estimate of analysis time | Optimal hardware utilization |
| Optimization | User | Selection of the best classifier based on user needs | Validated, deployable bioinformatics |
|  |  | GUI-enabled bioinformatic analysis |  |
|  |  | Selection of optimal pipeline at large scale, and deployment on laptop |  |

### Need and existing work

The difficulty inherent to the rapid and accurate classification of metagenomic sequencing data is a problem many researchers have attempted to solve, each publishing their own classification tools and comparing their performance against peer algorithms and/or third-party datasets.^2^ New algorithms are introduced on a constant basis, while older ones are updated to attempt to remain relevant. When you couple this with the steady progression of some sequencing platforms, it can be difficult for an end user to make an educated decision regarding the optimal sequencer/classifier pairing. META was developed to help users select from an ever-evolving landscape of emerging metagenomic classifier tools.^1^ As far as the authors can tell, only two other platforms have the potential to exist in the same application space as META: Galaxy and the Open-Community Profiling Assessment tooL (OPAL). However, while Galaxy could provide the foundation for side-by-side comparisons, there are no integrated evaluation tools^3^, and while OPAL provides more composition-level metrics for performance comparison, it lacks interactive data visualizations or table filtering options for classifier output^4^. Additionally, resource metrics such as run times and peak memory usage are not readily reported in the native installation state of OPAL, which requires a separate step of converting it to a Biobox Docker image. META provides all of these utilities by default, comparing the performance of more than ten classifiers and utilizing read simulators for the Illumina (MiSeq and iSeq instruments) and Oxford Nanopore Technologies (R9 and FLG flowcells) sequencing platforms^5,6^. META is openly available and can be accessed at the following github URLs: https://github.com/JHUAPL/meta-system, and https://github.com/JHUAPL/meta-simulator.

### Simulated versus real world sequencing modes

The META bioinformatics analysis pipeline enables the direct and simultaneous comparison of metagenomic abundance profiles derived from multiple classifiers using a set of *in silico* (simulated), or ‘real-world’ sequencing reads (Figure 1). In this way, the system supports two modes of classifier comparisons. ‘Mode 1’ enables the multi-classifier analysis of an *in silico* generated read set using sequencing platform simulators that produce data reflecting a user-defined abundance profile. Results are reported using three metrics: area under the precision recall curve (AUPRC) ^2^, the Euclidean distance between the user-defined abundance profile and the predicted abundance profile (L2) ^2^, and the metascore, which is a normalized combination of these two values. The AUPRC is a measure of how well individual taxa are identified and is more sensitive to low-abundance taxa, while the L2 metric is a measure of how reliably the abundance is calculated, meaning it is more sensitive to high abundance taxa. A complete description of the L2, AUPRC, and metascore can be found in the materials and methods.

**Figure 1.**
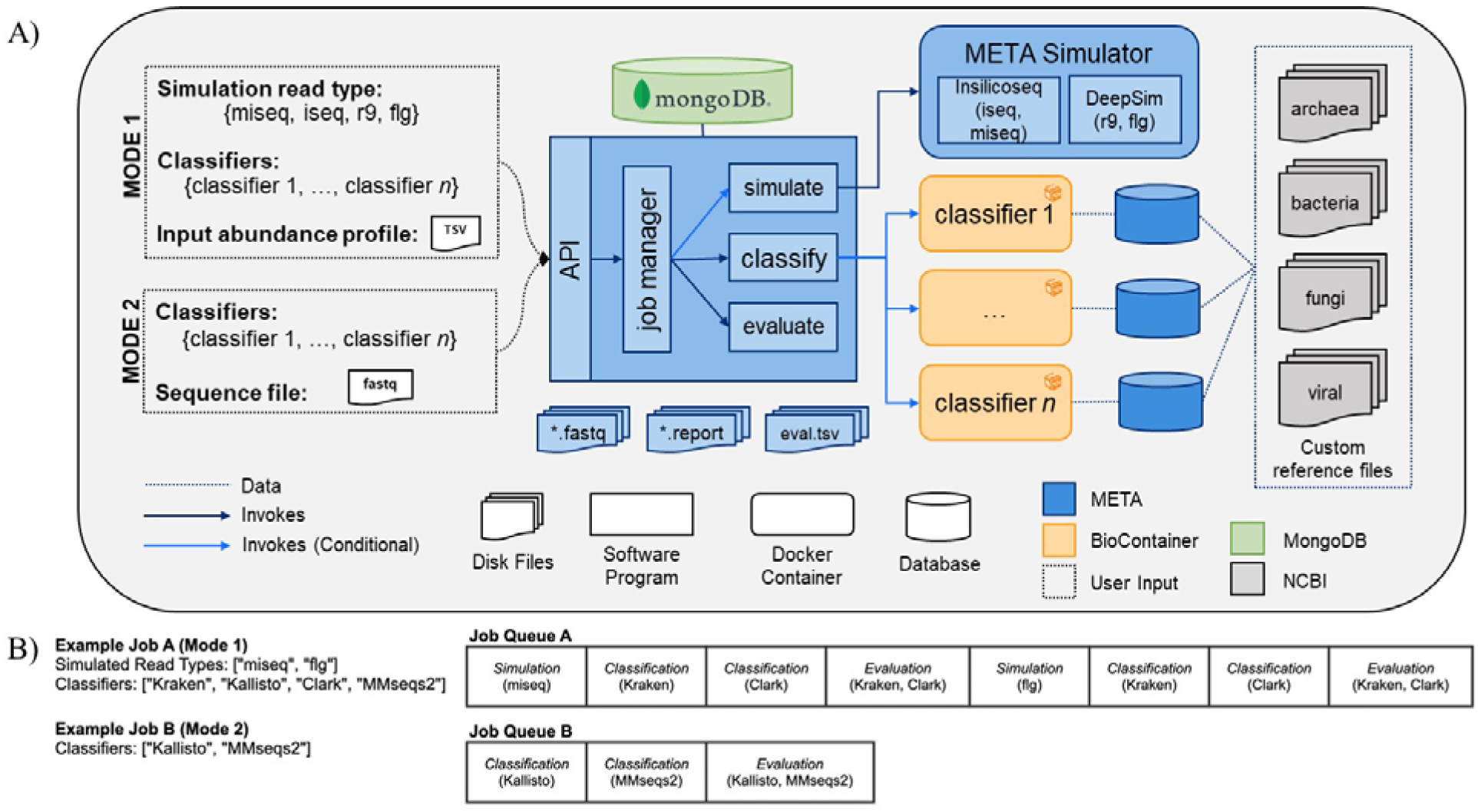
(A) META system architecture. The META system supports two evaluation modes: Mode 1 (*in silico* generated reads), and Mode 2 (real-world generated reads). Mode 1 enables classifier output to be compared to “ground truth” abundance profiles as supplied by the user. Reads can be generated using multiple sequencing platform simulators. After selecting which classifiers to evaluate and submitting a job, the system metadata is tracked in a MongoDB database while each selected classifier is run in series. (B) Sample META analytics workflows. Workflows are generated upon user request, and include serially-run simulation, classification, and evaluation modules.

‘Mode 2’ enables a comparison of output from multiple classifiers when the input consists of any FASTQ file, including those generated from real wet-lab experiments. Used in tandem, these two modes can help select the best sequencing and analysis approach prior to, and following, the experimental design phase.

### Classifiers, modularity, and standardization

The initial release of META contains a total of twelve classifiers (Table 2). There are seven k-mer-based algorithms: Bracken^9^, CLARK^10^, Kraken^11^, Kraken2^12^, KrakenUniq^13^, Mash^14^, and MMseqs2^15^. The remaining five classifiers use alignment-based algorithms: Centrifuge^16^, HS-BLASTN^17^, Kallisto^18^ DIAMOND^19^, and Kaiju^20^. The former three utilize nucleotide sequence databases, while the latter two utilize protein sequence databases.

**Table 2.** Metagenomic classification tools and associated metrics that are available in initial META public release. Custom database build size refers to the custom databases generated from the normalized set of references (units are gigabytes (GB) unless otherwise noted). Average peak memory usage and run times were derived from a total of 16 Mode 1 jobs of varying metagenomic composition (FASTQ file size averaging approximately 1.0 GB), and using the custom built databases associated with each classifier.

| No. | Classifier | Synopsis | Custom Database Build Size (GB) | Average Peak Memory Usage (GB) | Average Runtime (s) |
| --- | --- | --- | --- | --- | --- |
| 1 | Bracken | k-mer based abundance estimation from raw reads | 72 KB | 0.210 | 0.04 |
| 2 | Centrifuge | alignment based on BWT and FM-indexing schemes | 30 | 2.717 | 14.38 |
| 3 | CLARK | k-mer based species or genus level classification | 255 | 9.901 | 0.04 |
| 4 | DIAMOND | alignment based, SW against protein database | 24 | 0.935 | 145.47 |
| 5 | HS-BLASTN | An accelerated MegaBLAST search tool | 274 | 28.282 | 74.08 |
| 6 | Kaiju | alignment based on BWT and FM-indexing of protein sequences | 26 | 2.672 | 14.63 |
| 7 | Kallisto | alignment based, pseudoalignment procedure | 15 | 2.524 | 0.05 |
| 8 | Kraken | k-mer based, exact alignment | 45 | 0.240 | 4.95 |
| 9 | Kraken2 | k-mer based, exact alignment, and translated search mode | 32 | 0.205 | 0.32 |
| 10 | KrakenUniq | k-mer based, exact alignment with smaller memory requirements | 819 | 0.245 | 71.05 |
| 11 | Mash | k-mer based, locality sensitive hashing | 16 KB | 0.158 | 0.64 |
| 12 | MMseqs2 | k-mer based filtering with ungapped then ungapped alignments | 651 | 0.373 | 0.07 |

META is modular, making extensive use of Docker architectures via Biocontainers, to facilitate rapid updating or addition of new classification tools as they become available^7,8^. The system also utilizes currently-available common standards from the Global Alliance for Genomics & Health (GA4GH) to improve interoperability, security, privacy, data visualization, and compatibility^7,21^. Using these community-developed standards allows the system to have immediate utility across a wide range of scenarios and use cases.

### Test cases and simulated versus real world evaluation

To test the utility of META, our group designed a set of two experiments using real world samples sequenced on multiple platforms (Figure 2A). Use case 1 was designed to represent an environmental surveillance scenario, in which an aerosol sample, containing the human pathogen *Bacillus anthracis* and two non-pathogenic near-neighbors, was collected in a background of seven common environmental bacteria (Figure 2A, B, C, and D). Use case 2 was designed to represent a sample collected from an animal infected with a potential human pathogen. *Vaccinia* virus, a simulant for smallpox, was used as the pathogen, and the chicken cells used to propogate the virus represented the infected animal (Figure 2A). All samples were sequenced on both Illumina and Oxford Nanopore platforms and *in silico* reads were generated on the same platforms. FASTA and FASTQ files for both wetlab and *in silico* simulated samples were then run through META’s classifier algorithms, and reports were generated describing performance.

**Figure 2.**
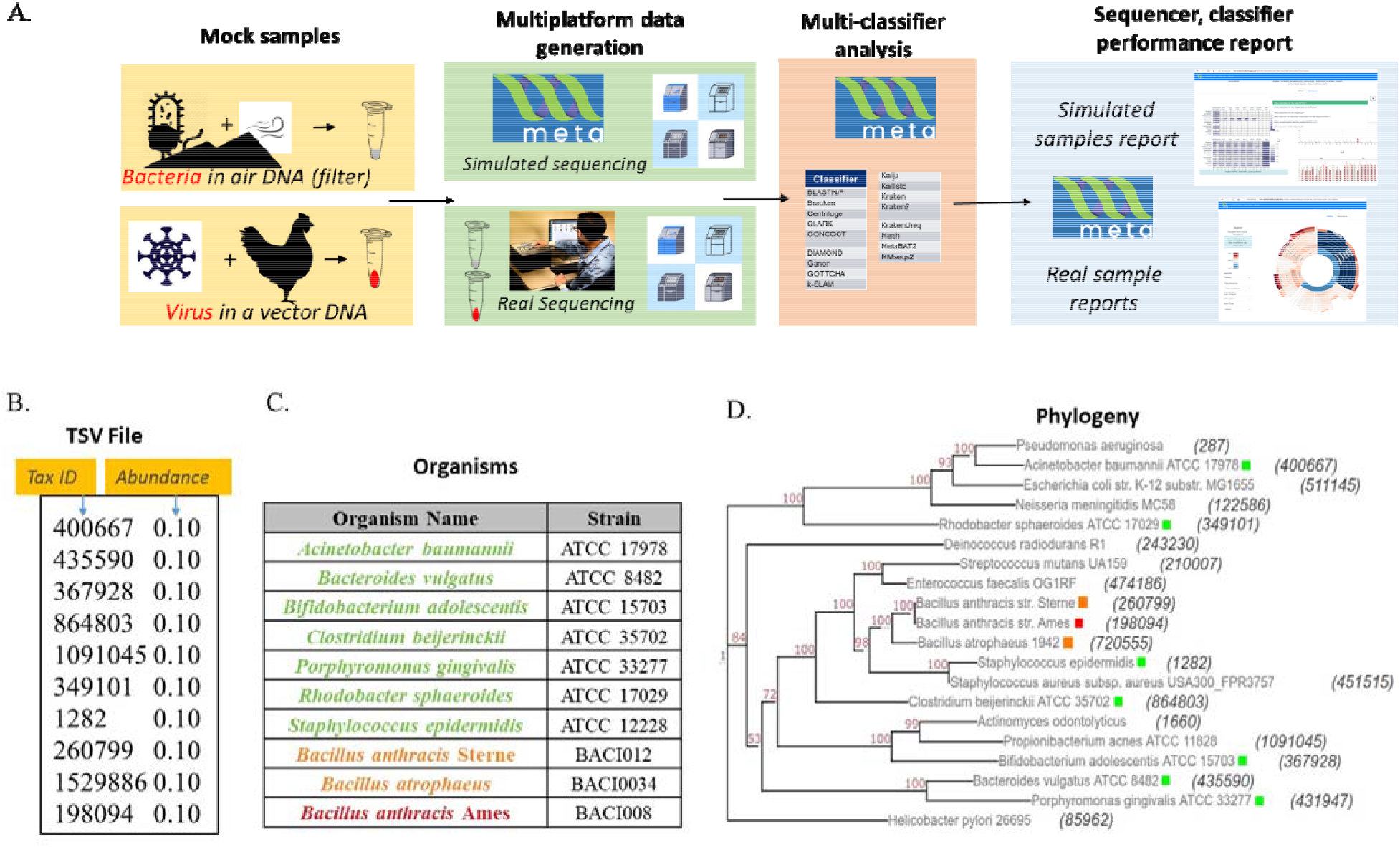
Sample use-cases test META capabilities. (A). Sequencing results from two mock metagenomic DNA communities are produced in the lab and mimicked in silico. Results using different classifiers are compared to the known sample content to identify the best performing sequencer/classifier combination. (B). Mode 1 user-generated TSV file for use case 1 mimicking a community found on an air filter. (C) Detailed components of community includes a threat agent (B. anthracis), two near neighbor organisms (B. anthracis Sterne and B. atropheus), and seven background environmental organisms. (D) Phylogenetic distance between each component of use case 1 community.

## RESULTS (usually contains a general description of the method followed by its validation)

### Survey and down-selection of classifiers for initial release

A literature search identified eighty-one metagenomic classifiers that were subsequently down-selected for inclusion in the initial release of the META system based on seven critical attributes (described in detail in Methods). Eighteen classifiers remained after down-selection criteria, and twelve were successfully implemented (Table 2). Classifiers used a range of algorithm strategies and custom reference databases.

### Use-Case 1

Mode 2 was utilized to evaluate the real-world data derived from experimental samples developed during use case generation. Classifiers were ranked by metascore for each read type (Illumina and ONT-generated reads) using the known spiking concentration for each component of the sample. Corresponding *in silico*-generated reads were then processed using Mode 1. Accurate strain and species identifications were prioritized in all analyses as this facilitated the differentiation of pathogens from background organisms.

The metascores for each sequencer/classifier pairing for use case 1 are shown in Figure 3. (L2 and AUPRC scores are displayed in Supplemental Figures S1 and S2). Separate analyses were run on 4 sample types: the pathogen alone, the pathogen and two near-neighbors, the seven background environmental bacteria, and all ten components together. Most classifiers scored similarly within each sample type and for each read type at the strain rank, with Mash having the lowest metascore in all cases. In general, the average metascore among all sequencer/classifiers pairings decreased as the total number of known organisms in a sample decreases. This is not unexpected, as the ability to discriminate between the total reads available become more difficult when the available reads in a sample are nearly identical. For the ‘mix of all 10’ organism samples, KrakenUnique achieved the highest metascore for the Illumina (MiSeq) read type, and Centrifuge the highest score for the ONT (R9 flowcell) read type.

**Figure 3.**
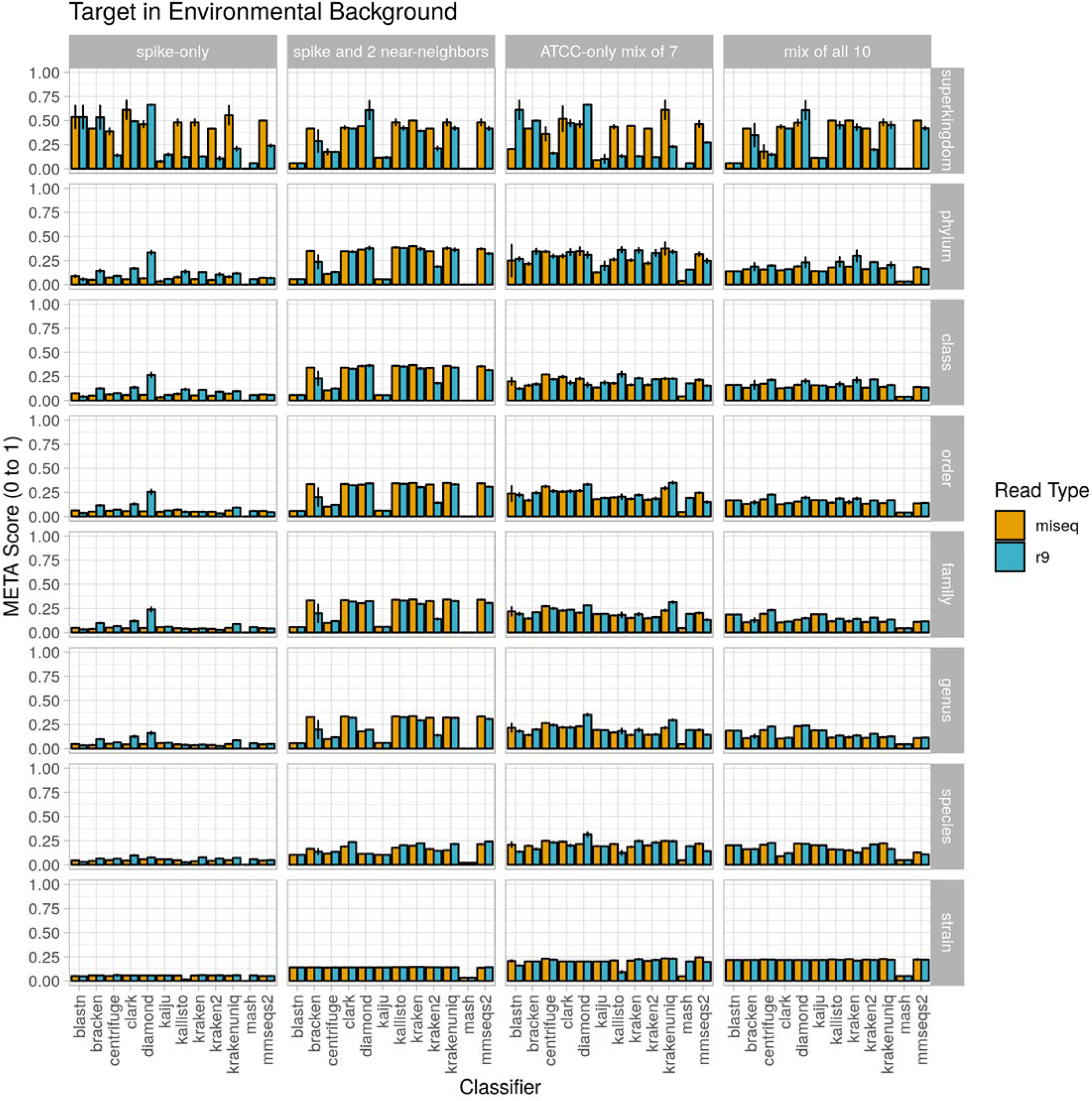
Use case 1 output. Classifier performance from *in vitro* data sets is illustrated using the metascore metric from superkingdom to strain taxonomic ranking. All twelve available classifiers were run using data generated from Illumina MiSeq and the ONT R9 flowcell. N=3.

Classifier performance for samples containing multiple organisms could also be determined by identifying (i) how many of the expected organisms were detected at any level, and (ii) how closely their respective abundance profiles matched the known spiking concentrations. While no classifier would be expected to identify all ten organisms, as the *Bacillus atrophaeus* subsp. *globigii* taxid was purposefully omitted in the custom reference database, Kraken, Kraken2, and HS-BLAST were the best performers, identifying all of the remaining 9 taxids when analyzing both Illumina and Nanopore reads (Table 3, columns two and three).

**Table 3.** Deviation of pathogenic target organism abundance from theoretical known value in use case 1. Values are representative of samples containing all 10 possible components at equal genome copy input. Nd= not detected.

|  | Number of Expected Taxids Identified in Output (Max = 9) |  | Calculated *B. anthracis* abundance (Theoretical Max = 10%) |  |
| --- | --- | --- | --- | --- |
|  | Illumina | ONT | Illumina | ONT |
| **Bracken** | 1 | 1 | nd | nd |
| **Centrifuge** | 5 | 4 | nd | nd |
| **CLARK** | 1 | 1 | nd | nd |
| **DIAMOND** | 1 | 1 | nd | nd |
| **HS-BLASTN** | 9 | 9 | 19.20 | 10.65 |
| **Kaiju** | 1 | 0 | nd | nd |
| **Kallisto** | 9 | 7 | 0.09 | nd |
| **Kraken** | 9 | 9 | 0.22 | 0.34 |
| **Kraken2** | 9 | 9 | 0.10 | 0.11 |
| **KrakenUniq** | 9 | 8 | nd | nd |
| **Mash** | 0 | 0 | nd | nd |
| **MMseqs2** | 9 | 8 | nd | nd |

The ability of the classifiers to identify the pathogenic component of the 10-organism samples was of particular interest. The three classifiers listed above successfully identified BA in Nanopore data sets, and the same three, plus Kallisto, identified BA from Illumina data sets. Theoretically, the *B. anthracis* read abundance should fall at approximately 10% for all of these sequencer/classifier pairs, but of the four classifiers that identified the pathogen at all using Illumina results, three were below 1%, while HS-BLASTN significantly over-represented its presence at >19%. The three classifiers identified the *B. anthracis* taxid from ONT results, with Two of the three classifiers that identified BA using Nanioire data sets significantly under-represented abundance (<1%) with only HS-BLASTN coming within 7% the expected value.

With respect to computer resource utilization, run times and peak memory usage varied widely based on the selected sequencer and classifier combination. Generally, though, no clear trend related these metrics with the metascore values, suggesting that users could not base their selection on these predictions alone. Full interactive reports are available for exploration on github (URL).

### Use case 2

Results from the second use case are displayed in Figure 4 (metascores) and Supplemental Figures S3 (AUPRC) and S4 (L2). For the Illumina data, the classifier achieving the highest metascore for “host”, “extraction1”, “spike1” and “spike2” was Mash (0.0572), CLARK (0.1787), CLARK (0.1165), and CLARK (0.0710), respectively. For the ONT data, the classifier achieving the highest metascore was DIAMOND (0.0571), CLARK (0.0861), DIAMOND (0.0673), and DIAMOND (0.0611), respectively (Figure 4). With respect to computer resource utilization, and only using those of the “spike2” sample, using the simulated Illumina read set CLARK was ranked 2^nd^ for runtime at only 0.42 seconds, however it was ranked 11^th^ for peak memory usage at over 118 GB RAM. Using the simulated ONT read set from the same sample profile, DIAMOND was ranked 11^th^ for runtime at 345 seconds, and 7^th^ for peak memory usage at only 8.6 GB RAM. Full interactive reports are available for exploration on github (URL).

**Figure 4.**
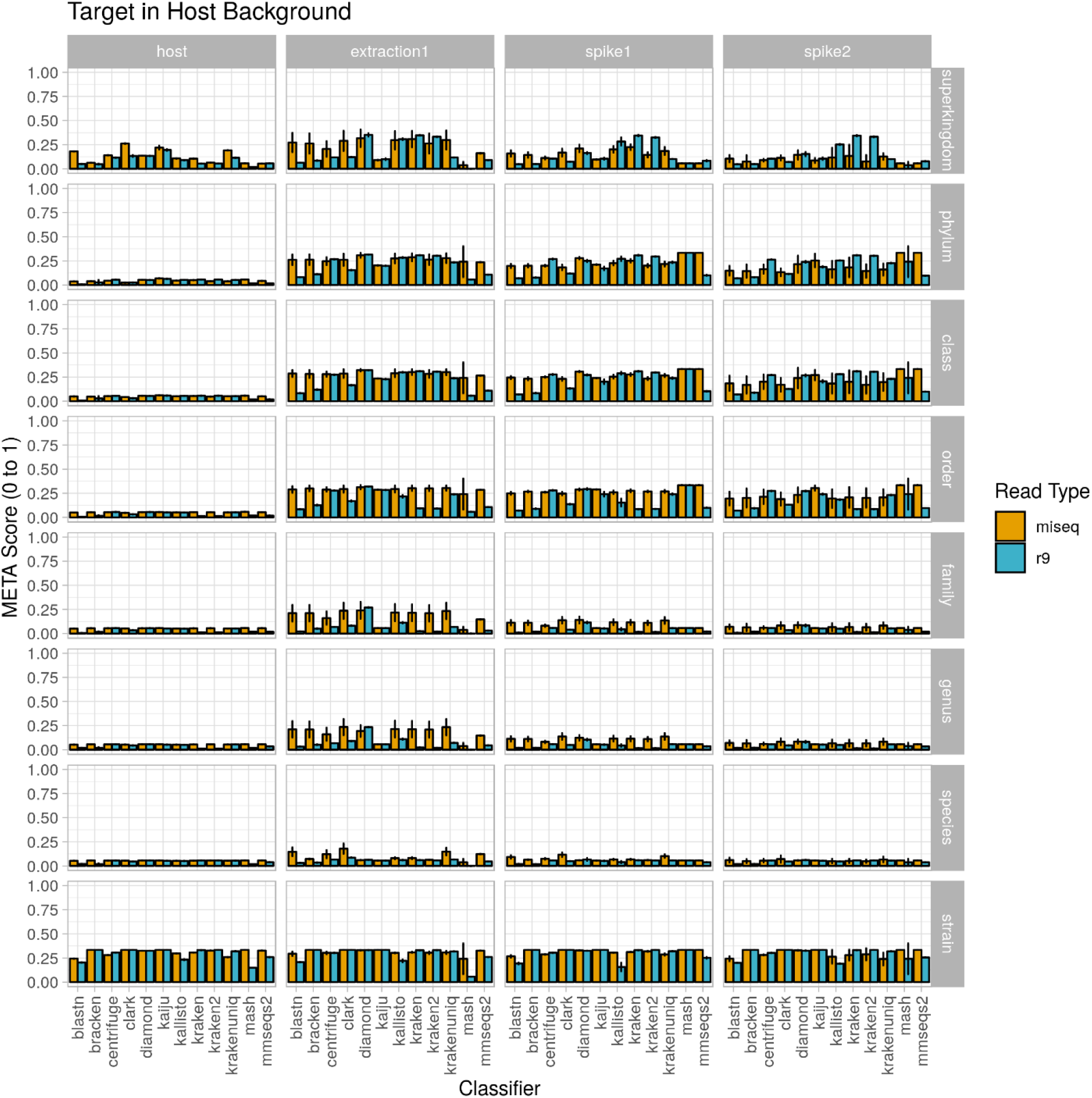
Use case 2 output. Classifier performance from *in vitro* data sets is illustrated using the metascore metric from superkingdom to strain taxonomic ranking. All twelve available classifiers were run using data generated from Illumina MiSeq and the ONT R9 flowcell. N=3.

A note here on the effects of this use-case (a single target of interest) on the value of AUPRC and L2. The AUPRC is very low across all taxonomic ranks for each sample type, implying there are many false positive classifications which is likely due to the fact that there is only a single ‘ground truth’ tax ID for these samples (the spiked organism). The L2 metric is much more variable across both taxonomic rank and sample type, and is actually closer to the ground truth profile at the stain rank compared to the higher ranks. This is somewhat counter-intuitive as one would assume that a more specific classification (e.g. strain) is more difficult than a less specific classification (e.g. species). However, and again since there is only a single ‘ground truth’ tax ID for these samples, even an abundance profile vector containing zeros for all tax IDs is closer to the ground truth than a vector that contains even a single tax ID that is not in the ground truth set at any significant abundance. These points coupled with the fact that L2 is in the denominator of the metascore calculation yields the higher strain rank scores seen in Figure 4.

Since the metascore is designed for the evaluation of classifier performance on more traditional metagenomic profiles, it was necessary to also check the deviation from the target ground truth abundance, as was done for the environmental experiment (Table 4). The target genome proportion reflects the target (*Vaccinia* Virus) to host (*Gallus gallus*) ratio in the sample type. The effective ground truth abundance is the abundance used for deviation calculations. This value for sample type (1) is zero, and all classifiers did not report it in their output abundance profiles, therefore their deviation is a perfect 0.00%. Sample types (2), (3), and (4) all have a target effective ground truth abundance of one, i.e. the input tsv column 3 for taxid 9031 (*Gallus gallus*) was set to 0, since it is not in the custom reference genome set. As the concentration of the target decreases, the deviation from target ground truth increases. CLARK was identified as the best performer using Illumina read type via metascore, and also consistently achieves the lowest deviation from target ground truth abundance for all sample types. Although DIAMOND was identified as the best performer in a majority of sample types using the ONT read type via metascore, it had a consistently high deviation from target ground truth abundance. Based on the deviation measure, CLARK, HS-BLASTN, or KrakenUniq may be a better substitute for identifying this target using the ONT read type.

**Table 4.** Deviation of pathogenic target organism abundance from theoretical known abundance in use case 2. Known abundance was set to 100%, as host genome was not included in the custom reference database. Nd= not detected.

|  | Host Only | Infected Lysate |  |  | Host Only | Infected Lysate |  |  |
| --- | --- | --- | --- | --- | --- | --- | --- | --- |
|  |  | Neat | 1:10 | 1:100 |  | Neat | 1:10 | 1:100 |
| Bracken | nd | -82.12% | -83.22% | -92.17% | nd | -84.44% | -94.30% | -99.36% |
| Centrifuge | nd | -4.80% | -36.93% | -85.23% | nd | -0.90% | -85.84% | -98.50% |
| CLARK | nd | -1.24% | -11.78% | -58.87% | nd | -0.70% | -66.78% | -95.15% |
| DIAMOND | nd | -94.42% | -94.74% | -95.22% | nd | -98.69% | -99.12% | -100.00% |
| HS-BLASTN | nd | -2.57% | -18.05% | -68.57% | nd | 0.00% | -77.48% | -97.32% |
| Kaiju | nd | -100.00% | -100.00% | -100.00% | nd | -100.00% | -100.00% | -100.00% |
| Kallisto | nd | -77.18% | -79.64% | -90.78% | nd | -82.05% | -90.82% | -100.00% |
| Kraken | nd | -81.57% | -84.82% | -92.93% | nd | -74.88% | -88.64% | -97.44% |
| Kraken2 | nd | -88.58% | -89.85% | -94.82% | nd | -86.11% | -94.44% | -99.37% |
| KrakenUniq | nd | -15.22% | -24.55% | -64.05% | nd | -1.40% | -69.38% | -95.80% |
| Mash | nd | -100.00% | -100.00% | -100.00% | nd | -100.00% | -100.00% | -100.00% |
| MMseqs2 | nd | -2.78% | -53.42% | -92.54% | nd | -37.80% | -88.07% | -99.18% |

## DISCUSSION

There is currently no bioinformatics tool that automates (i) the *de novo* simulation of Illumina and Oxford Nanopre Technologies (ONT) metagenomic sequencing data, (ii) the simultaneous classification of this *in silico* or real-world sequencing data using multiple algorithms, and (iii) provides performance metric information for each algorithm. Here, we present a software package and user interface called META to fill this gap. With META, researchers, can rapidly and inexpensively identify the best classification tool for a specific metagenomics use-case instead of settling on a strategy based on more subjective or qualitative comparisons potentially derived from unrelated data sets. META analysis outputs include a novel metric, the metascore, that utilizes both AUPRC to L2 measurements, allowing for a balanced and objective view of classifier performance. Feature-rich data visualizations are also produced, enabling the user to explore each analysis to the depth that suits their unique needs, eventually coming to their own conclusion about which classifier and read type combination will be most effective. The system is set up to rapidly add new classifiers, or update existing ones, so that a user can determine which sequencer and classifier combination is best suited to their use case, even as the technology continues to change and improve. Applications for this technology are numerous, including environmental monitoring, diagnostics, research, forensics, and industrial processes.

The next major release of META is expected to include a new mode that will enable reads from a known *in vitro* metagenomic sample dataset to be compared with an expected abundance profile. Assembly and alignment modules will also be added in order to support classification tools that require a contig or alignment file as input, respectively.

## METHODS

### Custom Reference Sequence Sets

In order to directly compare the performance of each classifier, a custom database was generated from the same set of reference organisms. This database included all archaea (347), bacteria (16,678), fungi (11), and viral (8,999) assemblies available on NCBI RefSeq that had a “latest” version status and an assembly classified as a complete genome. The approximate total size of the custom nucleic acid database was 69 GB, and the protein sequence database was 24 GB.

### Classifier Inclusion Criteria

Seven critical features were analyzed for each metagenomic classifier in order to down-select the final list for inclusion in the initial release of META: open source accessibility, availability on the Bioconda software package manager, date of last update, local data storage options, custom database input options, classification strategy, and analysis type. To be included in the META system, a classifier had to be be open source and available on Bioconda, a software package manager whose support increases ease of deployment and integration of the classifier. The presence of a classifier on Bioconda also implies that the tool is well-accepted and rigorously tested in the broader bioinformatics community. Recent updates were required to ensure that only actively maintained classifiers were selected, and as META requires classifiers to be run locally, any tool deployed only in a cloud-based environment was filtered out.

As mentioned above, in order to directly compare performance across classifiers, each tool had to allow for the use of a custom database, and due to the limited specificity of classifications based on marker genes, particularly 16S rRNA, all classifiers utilizing this strategy were excluded. This study aimed to review metagenomic classifier tools only, meaning other sequencing data analysis tool (e.g. assemblers, aligners, wrappers, etc.) were excluded, but could be included in future iterations of META in order to test and evaluate more complex analysis pipelines.

### System Architecture and Workflow

All META files and installation instructions are openly available on Github at https://github.com/JHUAPL/meta-system, and https://github.com/JHUAPL/meta-simulator. META is designed to run as a local web server based on Python Flask, accessible via RESTful API or user interface optimized for Google Chrome and Firefox using the Vue.js framework^22,23^(Figure 1A). The architecture leverages Bioconda and BioContainers, Docker container versions of the most commonly used metagenomic classifiers, in order to provide consistent deployment^7,8,24,25^. META has been designed to easily incorporate new metagenomic classifiers, providing a modular template for integration via a standard META YAML description^26^.

Towards interoperability, it was necessary to identify tool-specific commands that executed the four basic stages of metagenomic classification: download, build, classify, and report. This allowed for simultaneous processing of the same dataset by multiple classifiers without requiring individual inputs for each tool selected by the user. The following assumptions were made about all tools integrated with META; the tool database relies on a set of reference nucleotide or protein sequences for classification, and the tool can perform an analysis that provides output that allows for relative abundance calculations to be produced. These assumptions led to the following definitions within the META ecosystem: (i) Download: the command that downloads the set of reference nucleotide or protein sequences. Depending on the usecase, the set of reference sequences may already be present on the host machine. This is likely the case for those seeking to build custom reference databases. Currently, this command is not automatically executed within the META system. (ii) Build: the command(s) that build the database indices and relevant files from the set of downloaded reference sequences. An example of this is ‘kraken-build --db kraken_db --build’. Currently, this command is not automatically executed within the META system. (iii) Classify: the command that runs classification of sequence input. Classification analysis can be performed using several algorithms, including, but not limited to, distance metrics, string comparisons, or expectation maximizations (EMs). The command ideally contains an argument that specifies an output directory or output file path. An example of this is ‘kraken --db $db --output $output $input’. (iv) Report: the command pro that formats outputs. An example of this is ‘kraken-report --db $db $output’. An additional command may be necessary to rename the formatted report to ‘<tool_name>.report’. Some tools may not bundle a reporting utility, in which case, the META report file will be generated in the classify stage, and no report command needs to be identified. Notice that a set of installation commands are not required. This highlights the advantage of using BioContainer Docker images for deployment, as it does not require end users to install the tool and its dependencies prior to using it.

When a user is ready to process data using META, they must first submit a request in Mode 1 or Mode 2. To run a comparison using Mode 1 (*in silico* generated reads), an abundance profile must be provided in the form of a 3-column tab-separated variables (TSV) file. This file includes (i) the taxonomic ID (taxID) of the organisms making up the metagenomic community that will be simulated, (ii) the relative abundance of the associated taxID (abundance must sum to a total of 1.000000), and (iii) whether the taxID should (1) or should not (0) be considered in the relative abundance evaluation calculations. Users may opt to disclude certain taxID for abundance profile generation if the reads map to background genomic material or organisms of low interest. All input taxIDs in this file should map to an organism with a reference genome in NCBI’s RefSeq database, and a job will terminate and return an error message if a reference genome cannot be found for one or more taxIDs in an input TSV file. To run a comparison using Mode 2, the user must only provide a FASTQ file. All other parameters available to the user may be selected via check boxes on the job submission page, which include available classifiers to run and, for Mode 1 only, which read type(s) will be simulated for the provided abundance profile.

A META analytics workflow is automatically generated upon user request. Workflow components include simulation, classification, and evaluation modules that are processed serially. In Mode 1, for every simulated read type, there is an associated simulation module, and for every metagenomic classifier selected, there is an associated classification module. In Mode 2, the simulation module is excluded. In all modes, every FASTQ file has an associated evaluation module for computing metrics across all selected classifiers. Example workflows can be seen in Figure 1B. By distilling the workflow to these three composable modules (simulate, classify, evaluate) and four command types (download, build, classify, report), META prevents the “decision paralysis” inherent to more complex workflow environments and description languages^27,28^.

When a job is submitted, the user may view currently running jobs, and may select or download completed job reports for viewing or additional analysis. Most tables, graphs, and charts available in the report are fully interactive, sortable, and tips are provided where relevant. An additional ‘quick answers’ menu, in the form of Frequently Asked Questions (FAQs) is available in order to guide the selection of classifier(s) based on a particular metric of interest. When a FAQ is selected, this function automatically adjusts the data outputs to answer that specific question. Examples of questions include: ‘Which species has the highest L2?’, or ‘Which species and classifier combination has the largest AUPRC?’ (Supplemental Figure 5).

### Evaluation and Visualization

In Mode 1, the presence of known user-defined abundance profiles allows for direct comparison of classifier performance to ground truth values. This comparison is based on the ratio of the area under the precision recall curve (AUPRC) and the Euclidean distance between the user-defined abundance profile and the predicted abundance profile (L2) for each tool. These metrics are utilized because they are complementary as AUPRC is more sensitive to low abundance taxa, while L2 is more sensitive to high abundance taxa^2^. For AUPRC, each point on the curve represents the precision and recall scores at a specific abundance threshold. Abundance thresholds from 0 to 1 are used to generate the full curve. is generated, and the area calculated. The L2 metric is based on the pairwise Euclidean distance between the ground truth and classifier output taxa abundance vectors. Finally, a META score (referred to as metascore) is calculated from AUPRC and L2 by the following formula:

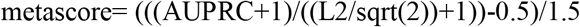

Dividing L2 by its maximum value (sqrt(2)) normalizes the range to that of AURPC (0 to 1). The addition of one to both metrics before taking their ratio bounds the ratio’s range from to 1/2 to 2. Finally, subtracting 0.5 from the ratio then dividing by 1.5 adjusts the range of the metascore to 0 to 1.

Classifier performance characteristics reported for both Mode 1 and 2 include a scatter/half-violin plot with a consensus call table, a parallel coordinates plot of resource utilization that includes CPU time, wall-clock time, and peak memory usage, and a sunburst plot of abundance that may be scaled and colored on various parameters to assist in data exploration. Visualizations were built using d3.js alongside the Vue.js frontend framework^23,29^. Each visualization is interactive and dynamic based upon user input and parameters attributed to a job. All tables available on the report page may be sorted and filtered and are linked to a single visualization.

The scatter/half-violin plot is designed to show the distribution of abundance calls across all classifiers for any number of read types (Mode 1 only) and specified ranks. Users may directly filter on any of these parameters within the plot by choosing from a dropdown menu or by zooming into a y-axis region. A table is also provided to display the specific point on the scatter plot that is attributed to a particular taxid upon cursor hover-over. An abundance thresholding distribution is provided to allow adjustment of the range of desired abundances to display.

Resource metrics including CPU time, wall-clock time, and peak memory usage for all selected classifiers (and read type for Mode 1) are plotted in a parallel coordinates plot. Each y-axis attributed to a metric is brushable to filter out undesired entries and is transferable (left-right) across the plot’s space for custom organization. Each line is hoverable to provide more information for that specific entry. A table is also provided that is directly linked to each line of the plot that is essential for identifying the best performing classifier-read type combination via metric sorting.

Finally, the sunburst provides a hierarchical representation of all taxids from superkingdom to strain ranks for a given classifier (and read type for Mode 1). Each slice size is based on the size of the abundance call for a given slice relative to the parent taxid and all sibling slices. Each slice is directly linked to an entry in a table and when clicked (table entry or slice), the plot dynamically updates to hide all ranks higher than the specified slice’s as well as all sibling slices at that rank. Color coding is selectable, and is based on either rank (Modes 1 and 2, default for Mode 2) or relative deviation from ground truth (input) abundance (mode 1 only, default for Mode 1; Supplemental Figure 6). Users may update the sunburst at any time with a dropdown selection of either the selected classifier and/or read type. An abundance threshold is also provided to allow users to adjust what range of reported abundances are to be observed in the plot, which is useful for classifiers that may call many unique taxids at exceedingly low abundances.

### Experimental Use-case Scenario 1: Pathogenic Target in Mixed Environmental Background

Use-case 1 was composed of equal genome copies from ten bacteria; *Bacillus anthracis* strain Ames (the “target” pathogenic organism), two taxonomic near neighbors of *B. anthracis*, and seven phylogentically separate environmental organisms that are also represented in the ATCC environmental microbiome standard set. Live *Bacillus anthracis* was acquired from BEI Resources (NR-411), while the near neighbors *Bacillus anthracis* strain Sterne UT238 and *Bacillus globigii* were grown from in-house stocks. All cells were streaked on TSA plates and incubated for 48 hours at 37°C. Colonies were selected from the plates and processed for nucleic acids using the DNeasy Blood and Tissue kit (Qiagen 69504). All other bacterial nucleic acids were obtained directly from ATCC as lyophilized reagents using the following catalog numbers: *Acinetobacter baumannii* (17978), *Bacteroides vulgatus* (8482), *Bifidobacterium adolescentis* (15703), *Clostridium beijerinckii* (35702), *Cutibacterium acnes* (11828), *Rhodobacter sphaeroides* (17029), *Staphylococcus epidermidis* (12228). Lyophilized nucleic acids were resuspended in nuclease-free water prior to combining. Organism nucleic acids were combined to generate four unique sample types for evaluation: (i) *B. anthracis* alone, (ii) *B. anthracis* and near-neighbors, (iii) 7 background organisms, and (iv) all ten components. For each sample type, all organisms were spiked at equal genome copy per organism with an expected metagenomic profile of equal relative abundances.

### Experimental Use-case Scenario 2: Pathogenic Target in Host Background

Use case 2 was composed of a viral pathogenic target present in a host cell background. Vaccinia virus strain MVA was acquired from BEI (NR-1) and was selected as the target viral pathogen due to ease of propagation, its DNA genome, and its similarity to the biothreat agent smallpox. VACV was grown in chicken (*Gallus gallus*) embryo fibroblast cells (ATCC CRL-1590) according to vendor recommendations. Briefly, cells were grown to 80% confluency at 37°C and 5% CO_2_ in growth media containing DMEM supplemented with 5% fetal bovine serum (FBS) and 5% tryptose phosphate broth, and infected at a multiplicity of infection (MOI) of 0.05 in inoculation media containing DMEM only. Following a 1-hour incubation period, inoculation media was removed and replaced with growth media. Three days post-infection, cells were scraped, centrifuged at 1200 x g for 10 minutes at 4°C, and resuspended in DMEM supplemented with 2% FBS. Cells were lysed using 3 freeze-thaw cycles and the final lysate was sonicated in ice water. Aliquots were stored at −80°C. Nucleic acid was isolated from both viral lysates and host cell lysates using the Qiagen QIAmp DNA Mini kit (51304). Viral and host nucleic acid was combined to generate four unique samples types: (i) host cells alone, (ii) neat viral lysate, (iii) 1:10 dilution of viral lysate in host background, and 4) 1:100 dilution of viral lysate in host background. Exact abundance of viral nucleic acid relative to host in samples 2-4 was not known prior to sequencing.

### Illumina and Oxford Nanopore Technologies Sequencing

All Oxford Nanopore Technologies (ONT) libraries were generated using Rapid barcoding sequencing kits (cat# SQK-RBK004). Sample libraries were multiplexed up to five samples per run and sequenced on three separate sequencing runs using ONT R9 flowcells (cat# FLO-MIN106D). Illumina MiSeq libraries were generated using Illumina Nextera XT library preparation kits (cat# FC-131-1096). All thirty libraries generated from both use cases were multiplexed together on a 2X300 paired end sequencing run using a 600 cycle MiSeq reagent kit (cat# MS-102-3003).

## ACKNOWLEDGEMENTS

Funding for this project was provided by the Defense Threat Reduction Agency (DTRA).

## AUTHOR CONTRIBUTIONS

RP integrated all project components, developed the analysis approach, and wrote large portions of the manuscript. AA led the software development team. BM developed the dynamic visualizations of META. LM and OA contributed to the development of the META system software, both front and back-ends. EF performed sample preparations, ddPCR, and lead the market survey. KV performed wet lab studies including sample preparation, DNA extractions, library preparation, and ONT and Illumina sequencing. BC contributed intellectually to the software design. SG contributed intellectually to the study and experimental designs and contributed to large portions of the manuscript. CB proposed and established the initial tool and study, provided project management and scientific direction, and contributed significant portions of the manuscript.

## DATA ACCESS

The META system is available on the github page of JHU/APL <URL>. Full installation of the META system includes running the open-source software from a Docker container; however, these docker containers are not part of the system itself (i.e. are not packaged with it before installation). Additional META software dependencies include Node 10+ and NPM 6+ (front-end), and Python 3.7+ (back-end). The software was developed on an Ubuntu 18.04 virtual machine containing 32 CPU cores, 512 GB RAM, and 10 TB disk space. Illumina and ONT data is available on NCBI’s SRA, accession PRNJXXXXX.

