## Supplementary figures and images for "The META tool optimizes metagenomic analyses across sequencing platforms and classifiers"

### Supplemental Figure 1

#### Slide 1
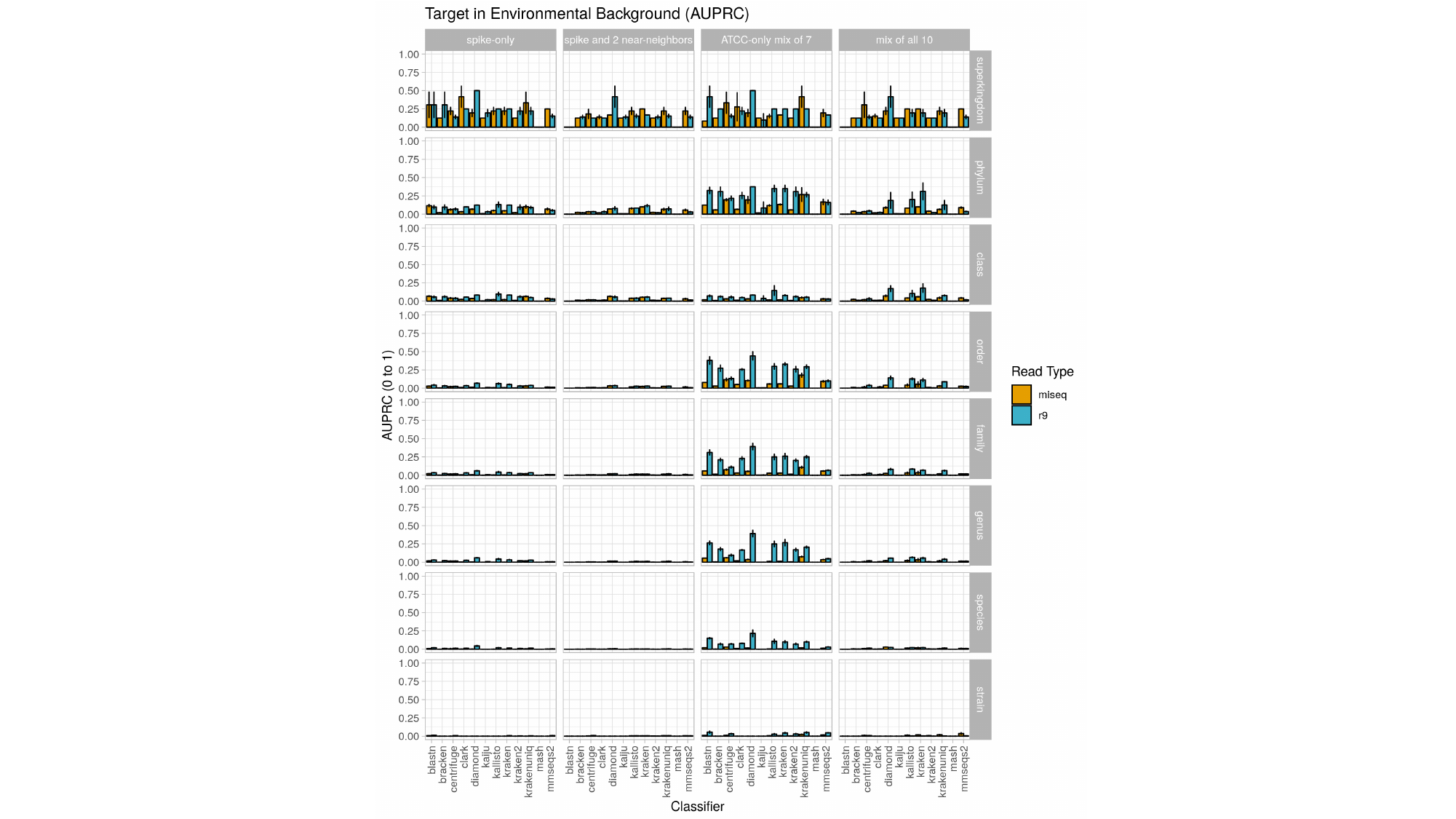

### Supplemental Figure 2

#### Slide 1
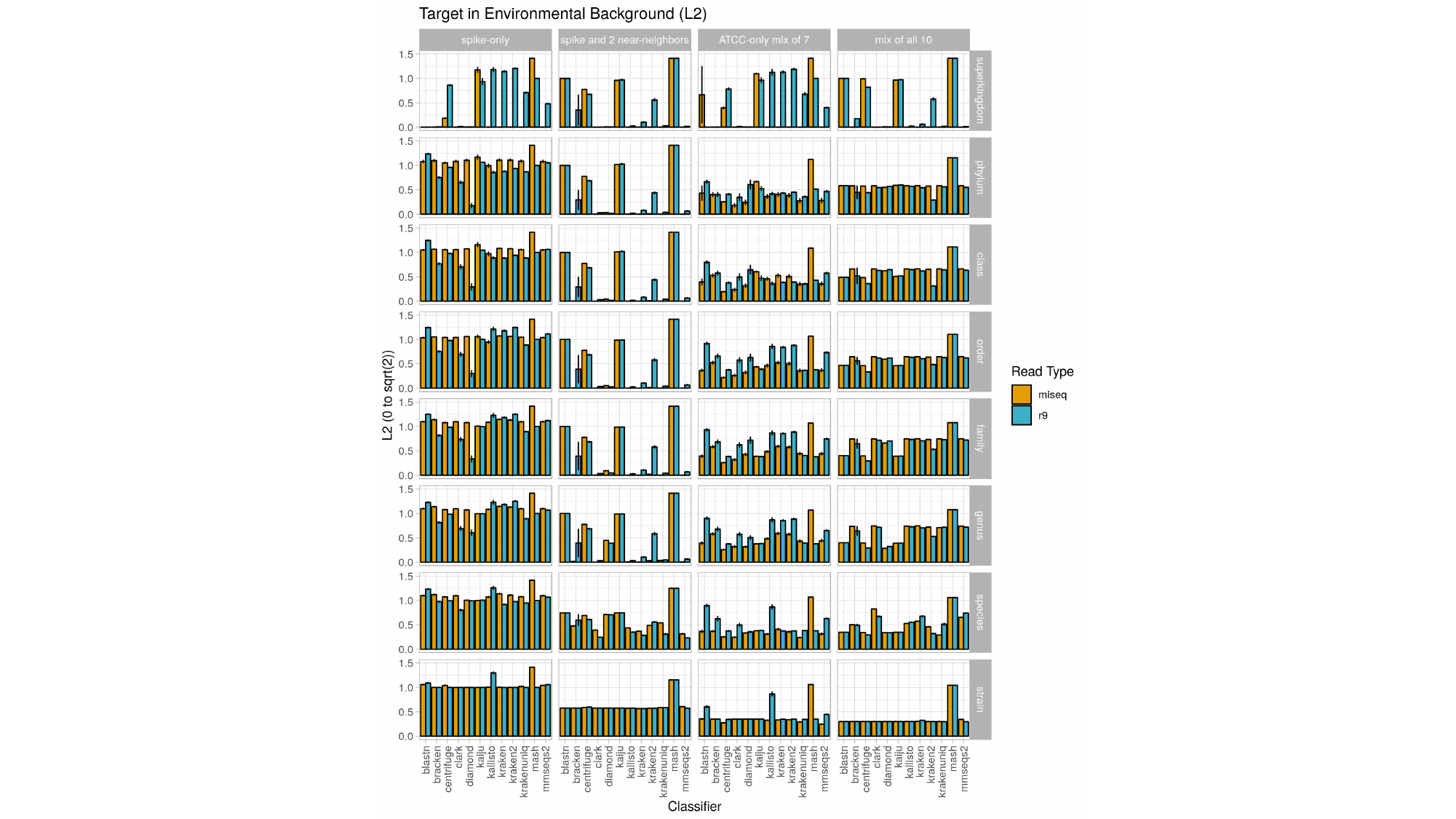

### Supplemental Figure 3

#### Slide 1
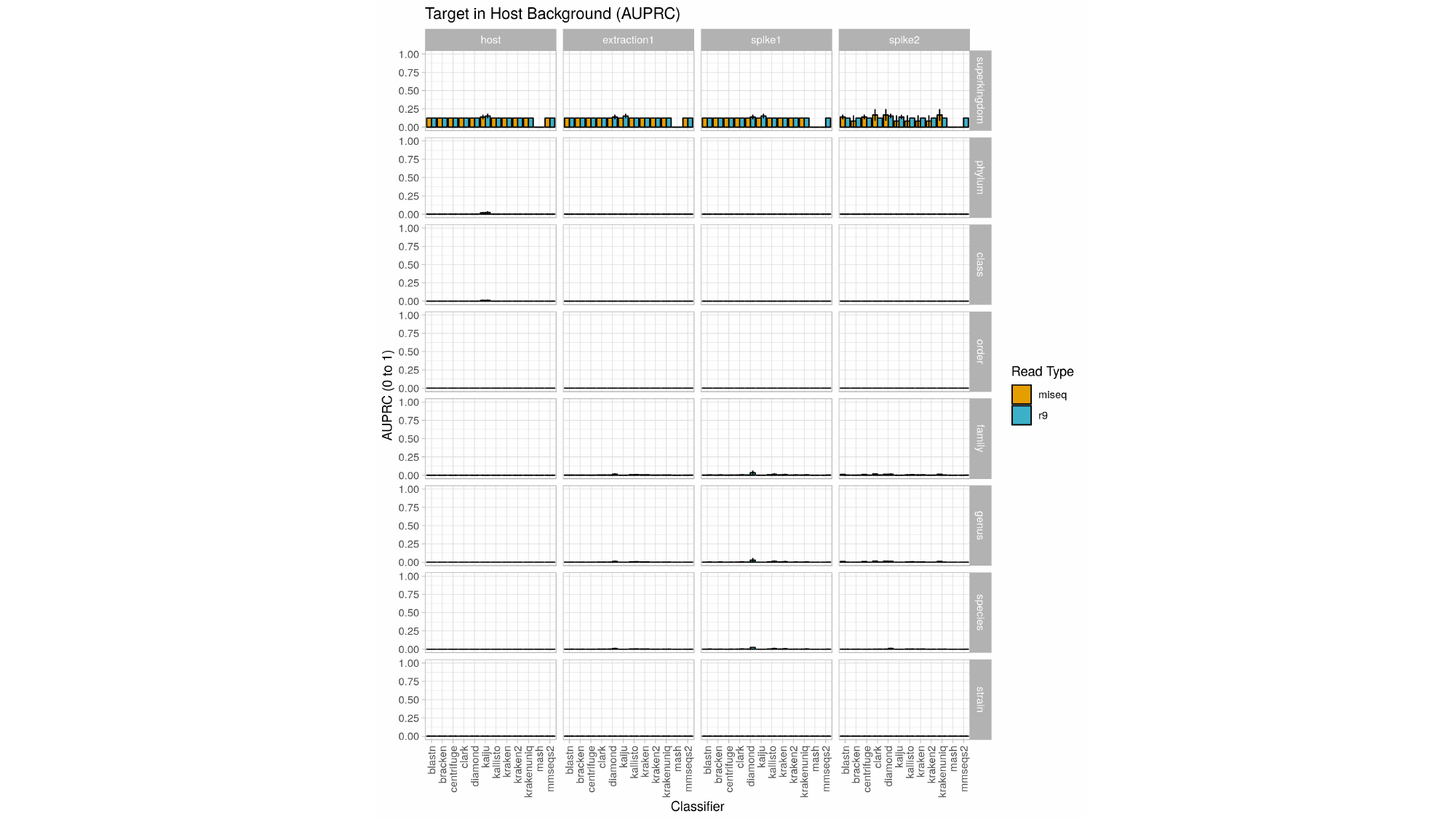

### Supplemental Figure 4

#### Slide 1
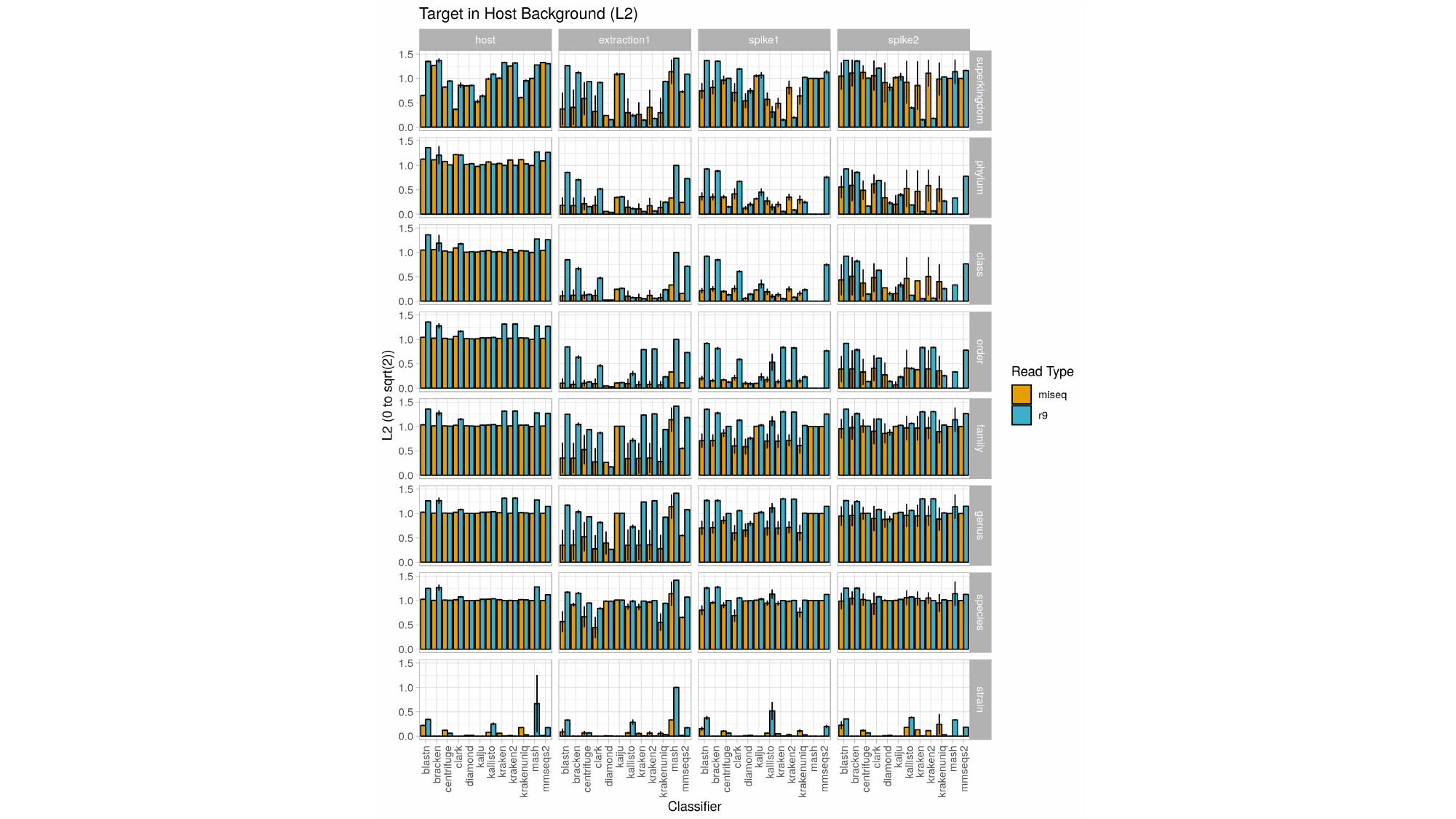

### Supplemental Figure 5

#### Slide 1
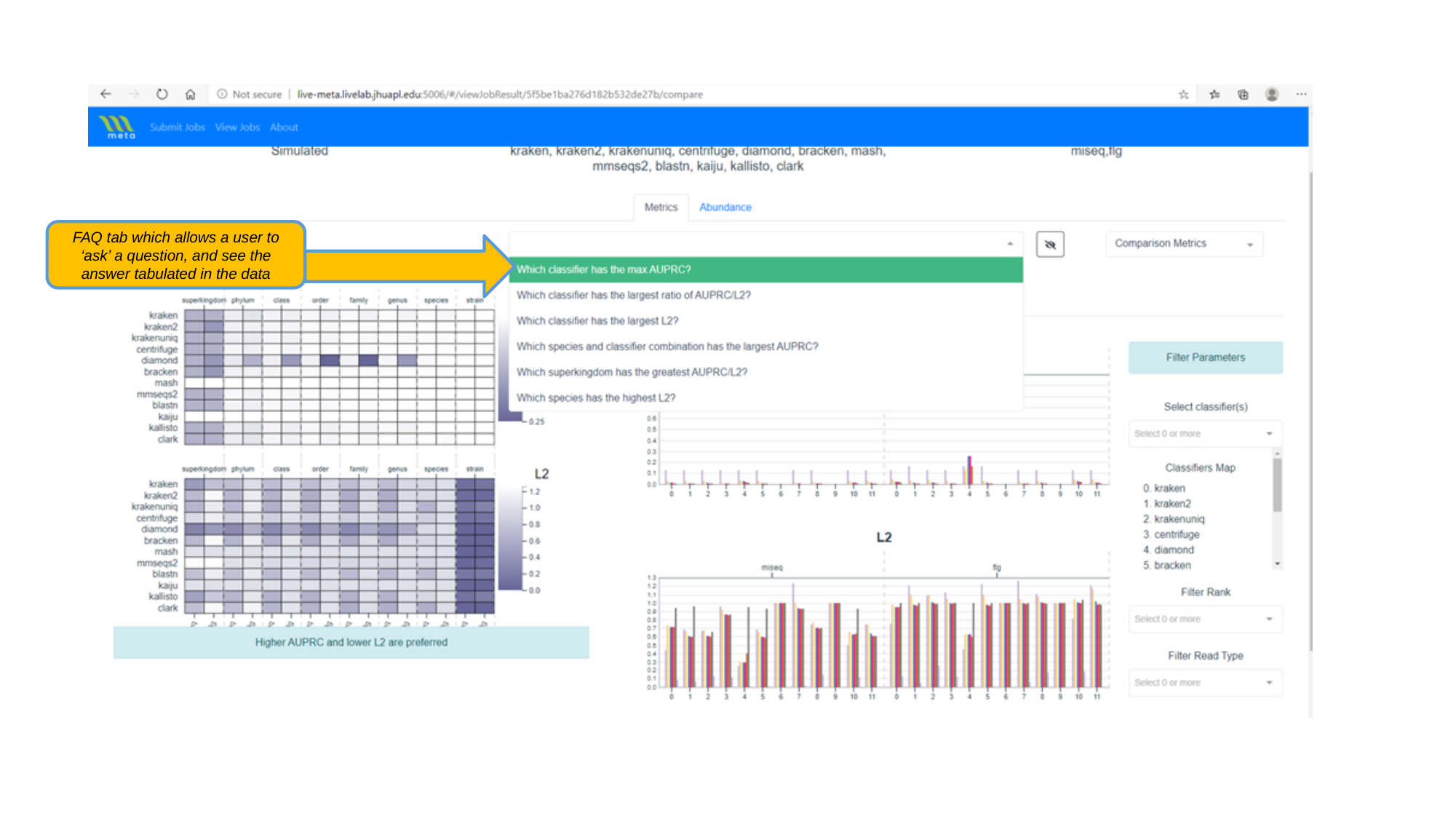

FAQ tab which allows a user to ‘ask’ a question, and see the answer tabulated in the data

### Supplemental Figure 6

#### Slide 1
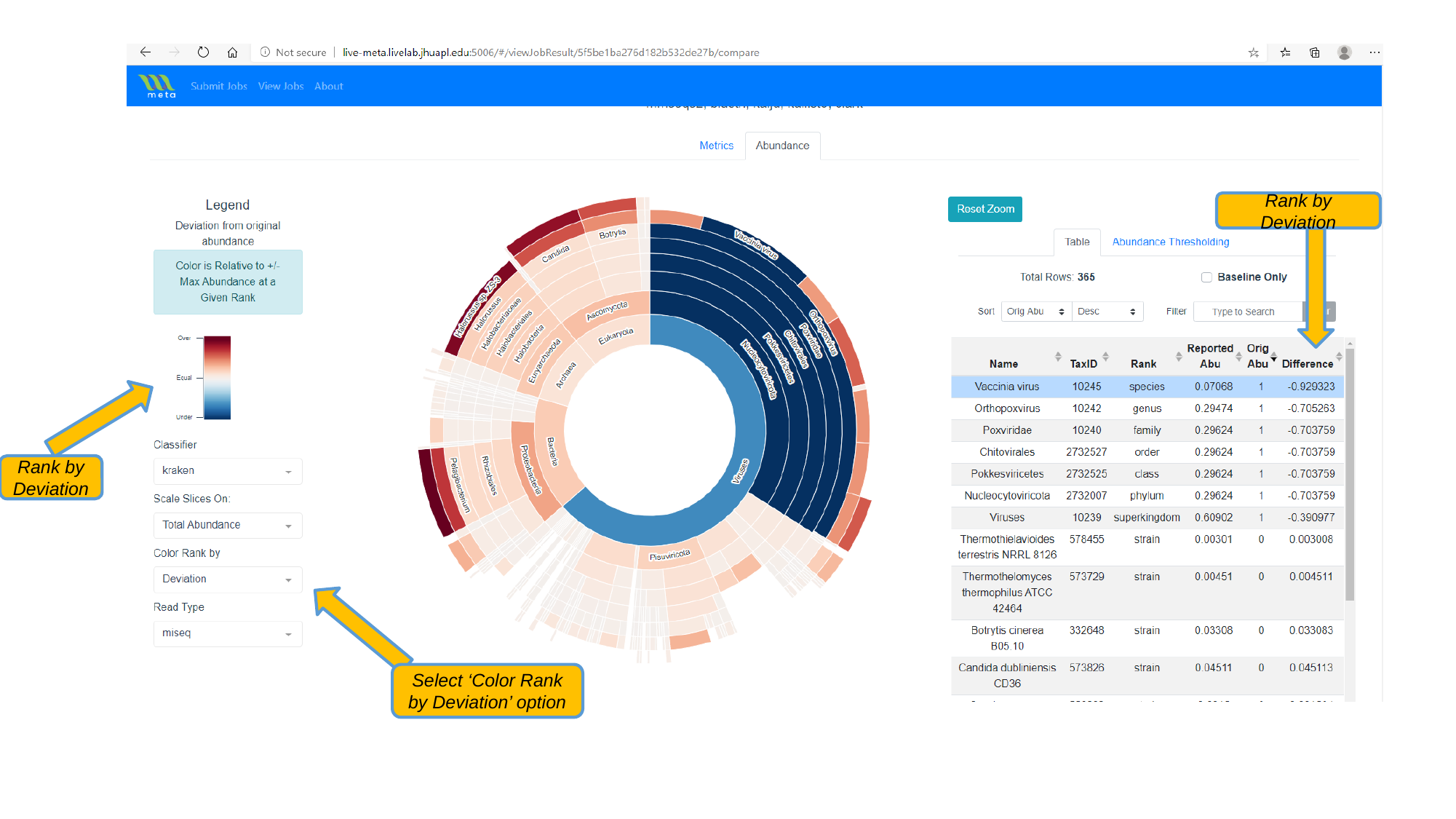

Rank by Deviation
Rank by Deviation
Select ‘Color Rank by Deviation’ option
